# A High-Fidelity 3D Fluid-Structure Interaction Framework for Predictive Microfluidic Design

**DOI:** 10.64898/2026.04.28.721227

**Authors:** Lingyue Shen, Yumiao Zhang, Yongxin Chen, Xu Ding, Pinjing Wen, Cheng Wang, Pengtao Sun, Shihua Gong, Jinchao Xu, Jiarui Han, Yan Chen

## Abstract

The commercial maturation of microfluidics remains bottlenecked by empirical prototyping and an absence of predictive digital design capabilities. Because optimizing advanced technologies such as passive particle separation fundamentally hinges on the precise coupling of fluid dynamics and particle mechanics, conventional two-dimensional or decoupled fluid simulations inherently fail to capture authentic multiscale behaviors. To bridge this gap, we establish a high-fidelity three-dimensional fluid-structure interaction framework combining a high-order Arbitrary Lagrangian-Eulerian mapping-based finite element method with a localized hierarchical dynamic mesh strategy. Engineered to accurately resolve complex multiscale hydrodynamics, this architecture utilizes deterministic lateral displacement structures as a stringent test case. Validated against experimental data for rigid microspheres and tumor cells, the framework predicts transport trajectories and critical separation diameters with sub-micron precision. Crucially, the simulation explicitly resolves the M-shaped spatial fluctuation of local size thresholds alongside the dynamic vertical migration of particles. Unveiling these hidden physical mechanisms provides a deterministic explanation for highly debated phenomena such as mixed-mode transport. By enabling the rigorous *in silico* evaluation of complex non-periodic architectures, this framework serves as a powerful instrument for predictive structural optimization. Such capabilities establish the essential infrastructure for microfluidic digital design, accelerating the transition from empirical trial-and-error to precision simulation-driven engineering.

## Introduction

Today’s artificial intelligence revolution, from large-scale language models to drug discovery, is built upon a foundation of high-performance processors. These critical hardware components, including CPUs, GPUs, and specialized ASICs, are themselves the product of a decades-long paradigm shift: from physical prototyping to simulation-driven digital design. The engine of this transformation is Electronic Design Automation (EDA), a sophisticated software ecosystem enabling engineers to design, verify, and optimize chips with billions of components *in silico*. Without EDA to manage this extreme complexity and mitigate the prohibitive costs of fabrication errors, the modern semiconductor industry and the digital world it supports would not exist.

In stark contrast, the field of microfluidics has yet to fully realize its transformative potential despite being long heralded as a parallel lab-on-a-chip revolution for biology and medicine. We argue that this persistent gap between promise and commercial reality stems from the critical absence of a dedicated Microfluidics EDA ecosystem. Consequently, the field remains shackled to a build-and-test paradigm characterized by a slow and inefficient cycle of iterative physical fabrication. This reliance persists because general-purpose Computational Fluid Dynamics (CFD) tools exhibit two fundamental shortcomings. First, they fail to resolve the extreme multi-scale disparity between centimeter-scale device footprints and micrometer-scale fine structures. Second, they lack the algorithmic efficiency to compute fully coupled fluid-structure interactions (FSI). Because advanced microfluidic technologies such as passive particle separation, inertial focusing, and deterministic particle manipulation depend heavily on the precise coupling of fluid dynamics and particle mechanics, optimizing these devices fundamentally hinges on accurately capturing this dynamic interplay. Traditional methods relying on decoupled fluid-only simulations or simplified two-dimensional (2D) models are therefore inherently incapable of predicting the authentic physical behaviors within such complex systems.

Deterministic Lateral Displacement (DLD) epitomizes this challenge. As a passive, label-free technology for high-resolution particle sorting, DLD exploits the interaction between particles and a tilted array of micropillars to separate cells based on size, shape, and deformability^1–4^. Its applications include sorting diverse microparticles such as parasites^5^, blood cells^6^, circulating tumor cells^7^, bacteria^8^, and spores^9^, and have more recently expanded to nanoscale exosomes^10^ and DNA isolation^11,12^. However, even for fundamental configurations involving rigid spheres and cylindrical obstacles, the intricate interplay between boundary-layer hydrodynamics and particle trajectories defies robust theoretical prediction, leaving a significant gap in our understanding of particle-mode behavior^13^. Thus, leveraging simulation tools to predict particle trajectories offers a critical pathway for identifying optimal structural designs. To achieve this, predictive design tools must overcome two fundamental and interdependent challenges. The first challenge is physical fidelity. A fully coupled FSI framework that can precisely capture the intricate dynamic fluid-structure interfaces is required to reliably quantify the local hydrodynamic perturbations introduced by the particles and the resulting forces acting upon themselves. The second challenge is computational efficiency. Accurately simulating micrometer-scale particles flowing through centimeter-scale devices with micrometer-level resolution requires prohibitive computational resources. This multi-scale complexity has historically necessitated the use of 2D approximations, which fundamentally fail to capture the three-dimensional (3D) velocity distributions required for precise trajectory prediction.

Various numerical strategies have been developed to tackle the complexities of FSI in DLD device, yet each maintains inherent trade-offs between physical fidelity and computational efficiency. Dissipative Particle Dynamics (DPD)^14,15^, a stochastic mesoscopic method, excels in capturing fluctuating hydrodynamics but often struggles with the high computational cost required to resolve macroscopic boundary conditions and sharp fluid-solid interfaces^16–18^. On the other hand, the Lattice Boltzmann Method (LBM)^19,20^ handles parallelization and complex geometries efficiently, but it typically requires high-resolution uniform grids to maintain stability, thus becomes computationally prohibitive for multi-scale systems where sub-micron boundary layers must be resolved within centimeter-scale domains^21–23^. To bypass the complexities of dynamic meshing, researchers often employ body-unfitted mesh methods like the Immersed Boundary Method (IBM)^24^ and the Fictitious Domain (FD)^25,26^ method. These techniques use Dirac Delta functions or Lagrange multipliers to impose kinematic constraints on a fixed background grid, but this detachment often reduces the accuracy of capturing steep velocity gradients near fluid-structure interfaces^21,27–29^. If researchers choose body-fitted meshes to achieve superior interfacial fidelity, the Arbitrary Lagrangian-Eulerian (ALE) method is the standard choice^30,31^. However, the large solid displacements inherent to particle separation induce severe mesh distortion, necessitating frequent and costly global remeshing that effectively precludes high-throughput screening^32–35^.

To overcome these challenges, our group previously established a robust class of monolithic interface-fitted/fictitious domain-finite element methods for 2D scenarios^36– 38^. While these methods successfully bypassed the need for global remeshing in 2D, extending them to realistic 3D domains remained computationally prohibitive. Transcending this dimensionality barrier, we developed a novel fluid-structure interaction framework engineered to accurately resolve complex multiscale hydrodynamics in microfluidic devices. Our framework achieves both physical fidelity and computational tractability through three key architectural innovations comprising a hierarchical mesh strategy, high-order spatiotemporal numerical schemes, and a localized dynamic mesh updating strategy. By integrating these approaches, the framework renders fully coupled 3D FSI simulation of entire DLD devices possible on limited computational resources for the first time. Validated against physical experiments utilizing both synthetic particles and biological cells, this virtual prototyping tool accurately predicts complex transport trajectories. Crucially, the framework enables a deep investigation into the fundamental physics of DLD and provides deterministic explanations for highly debated phenomena such as mixed-mode transport. Illuminating these hidden hydrodynamic mechanisms empowers engineers to systematically optimize device performance *in silico* prior to physical fabrication. Built upon universal physical principles, this robust infrastructure remains broadly applicable to diverse microfluidic systems including cell sorting, hydrodynamic focusing and trajectory modulation. Ultimately, this work establishes a definitive computational pathway for predictive device optimization. By shifting the paradigm from empirical trial-and-error to high-fidelity virtual prototyping, this computational infrastructure will fundamentally overcome traditional hardware development bottlenecks and accelerate the discovery of next-generation microfluidic technologies.

## Results

### Principle of the 3D FSI simulation framework

We developed a high-precision, high-efficiency FSI computational framework capable of simulating coupled fluid-structure motion in complex 3D domains. As a test case for this framework, we modeled a DLD device composed of a large-scale 50 × 1000 array of periodically arranged micropillars and simulated the resulting particle trajectories (Fig. 1a). Utilizing this 3D FSI model, we sequentially calculated the flow field, constructed the computational mesh, initialized particles of specific sizes at designated starting coordinates, and computed their continuous dynamic trajectories.

**Figure 1.**
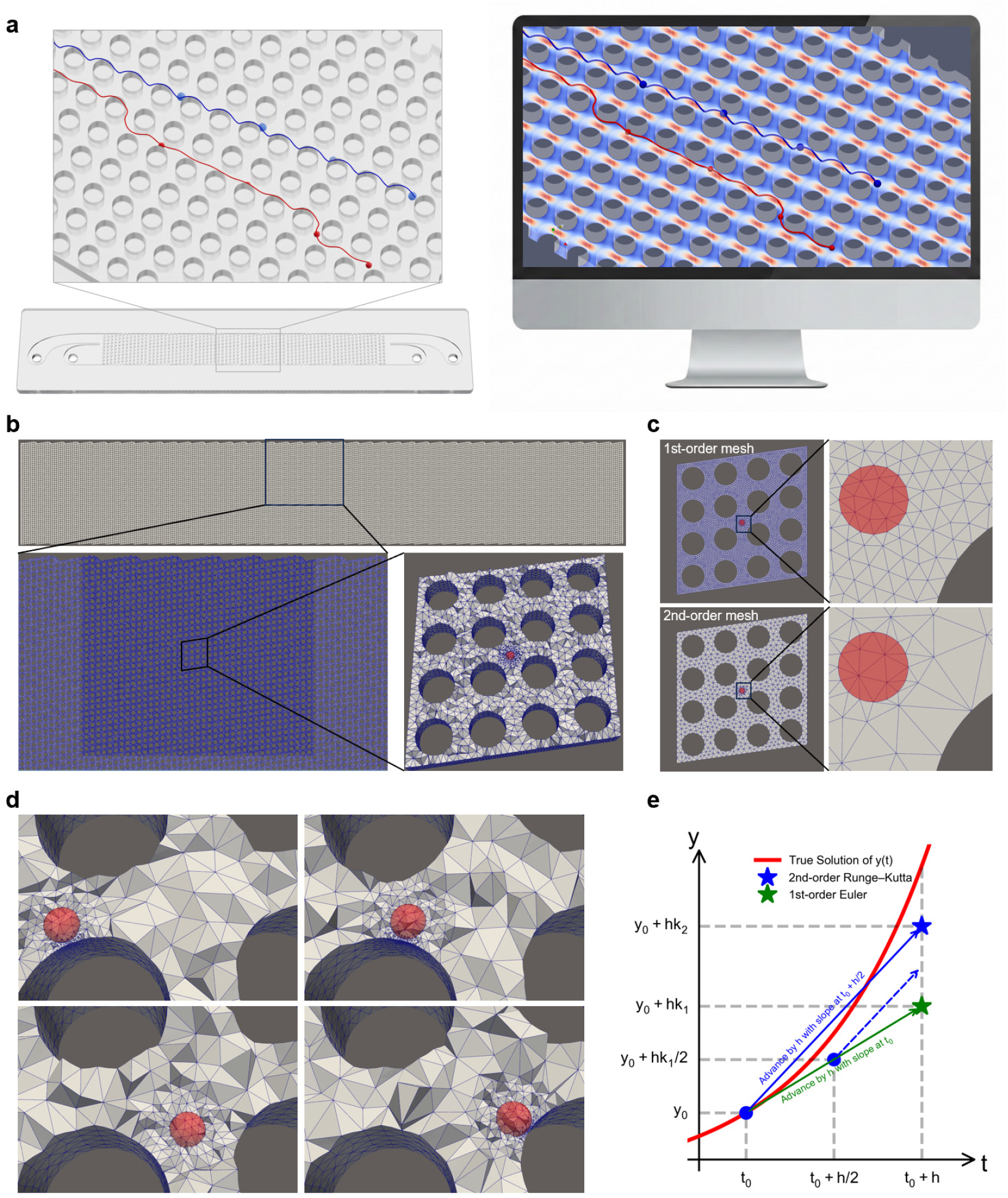
Schematic illustration of the 3D fluid-structure interaction (FSI) simulation framework. (**a**) Comparison of experimental and simulated particle dynamics, demonstrating excellent agreement between the trajectories of differently sized particles within a DLD device and their corresponding *in silico* predictions. (**b**) The hierarchical mesh strategy. (**c**) First-order mesh and second-order isoparametric mesh with same discretization error. (**d**) The dynamic adaptive mesh technique, demonstrating precise and continuous tracking of the moving fluid-solid interface. (**e**) Temporal discretization schemes, contrasting the first-order Euler method with the second-order Runge-Kutta integration.

Conventional simulation algorithms typically rely on periodic boundary conditions to approximate flow within a single unit cell. Our framework overcomes this limitation by directly simulating large-scale arrays without assuming structural periodicity. This capability is essential for analyzing complex designs where pillar geometry and spacing vary spatially. Furthermore, our framework employs a fully coupled solution strategy to resolve complex fluid-structure interactions. Given that precise particle tracking inherently requires time-dependent evolution, our framework utilizes high-order spatiotemporal discretization schemes to predict these dynamic behaviors across the entire microfluidic device.

To address the multi-scale challenges of FSI, the framework introduces a hierarchical mesh strategy. The computational domain is partitioned into three nested regions of increasing resolution: a coarse outer mesh for the flow field of a large area of pillar array, an intermediate mesh for the pillar array of interest, and finally an ultra-fine, dynamically moving mesh that envelopes each particle (Fig. 1b). The fully coupled fluid-structure solver is applied only in the vicinity of the moving particle. Specifically, the solution fields of pure fluid flow derived from the outer region establishes the boundary conditions for the intermediate region, which subsequently provides the highly resolved boundary conditions for the innermost region. This cascading approach ensures that the steep velocity gradients in the boundary layer near pillars can be resolved with sub-micron precision, and sophisticated FSI effects around moving particles can be achieved locally with higher precision as well, while the computational load remains minimal elsewhere. Consequently, compared to traditional uniform meshing techniques, this hierarchical mesh generation strategy accelerates computation speed by two orders of magnitude while maintaining the same numerical accuracy. Furthermore, the methodology is general and can be applied to any device geometry or boundary condition, enabling simulation of a very large domain without requiring periodic structural assumptions.

We employ high-order discretization schemes in both space and time to greatly improve computational efficiency while preserving numerical accuracy. For spatial discretization, second-order isoparametric finite elements (body-fitted to the fluid-structure interfaces) are used, and the mesh is adaptively refined in critical regions to achieve high resolution where needed. This high-order spatial scheme captures fine interface details with fewer mesh elements compared with the first-order finite element mesh, thereby controlling the overall mesh size and cost (Fig. 1c). For temporal discretization, we use a second-order time-integration scheme (a second-order Runge-Kutta method), which provides greater accuracy and stability than a first-order scheme and permits the use of larger time step sizes (Fig. 1e). Together, compared to a conventional first-order baseline, these high-order strategies improve the computational speed by three orders of magnitude without sacrificing solution accuracy.

To accurately resolve the coupled dynamics of FSI, we employ a dynamic mesh technique. Unlike static grids, the computational mesh continuously deforms and moves with the solid particle, thereby enabling precise tracking of the fluid-structure interface (Fig. 1d). At each time step, the mesh node positions are updated according to the posterior mesh velocity field driven by the structural velocity to ensure that the mesh follows the structural trajectory. This approach effectively accommodates large displacements without the need for computationally expensive global remeshing. Remeshing for the innermost fine FSI region is performed only when severe fluid mesh deformation leads to unacceptable mesh quality. Consequently, high-fidelity resolution of the fluid-structure interface is consistently maintained throughout the simulation. Compared to uniform meshes utilizing first-order schemes, this framework achieves a five-order-of-magnitude acceleration in overall computational speed while maintaining the same numerical accuracy. The successful application of this combined hierarchical and dynamic meshing approach to simulate comprehensive 3D particulate fluid and particle trajectories within a 3D DLD chip domain is visually demonstrated in Supplementary Movie 1.

### Depth-dependent separation thresholds and dynamic longitudinal migration

To evaluate the intricate 3D transport dynamics within DLD arrays, we utilized our fully coupled FSI framework to simulate particles navigating the microfluidic channel. The separation performance of these devices was fundamentally governed by the critical diameter (*D*_*c*_), which served as the specific size threshold dictating whether a particle followed a zigzag path or lateral displacement. Despite this straightforward definition, physical experiments frequently revealed a transitional mixed-mode interval where intermediate-sized particles exhibited alternating transport behaviors. Our high-fidelity simulations uncovered the physical origins of this phenomenon by revealing two previously obscured depth-dependent mechanisms.

First, the simulations captured a pronounced dynamic longitudinal migration of particles along the vertical z-axis. As particles traversed the micropillar array along the primary flow direction, they progressively migrated away from the unstable channel midplane. Different particle sizes displayed distinct vertical trajectory characteristics during this process. Tracking a smaller 7 µm particle in the zigzag mode revealed cyclic vertical fluctuations that perfectly corroborated experimental observations of periodic particle defocusing around pillars (Fig. 2c). Conversely, a larger 9 µm particle in the displacement mode continuously migrated downwards until it ran adjacent to the bottom wall (Fig. 2d). An intermediate 7.8 µm particle demonstrated a dynamic transition, beginning with a zigzag step before permanently switching to the displacement mode (Fig. 2e). Statistically, after traversing the entire 36 mm device, 7 µm and 9 µm particles initially distributed uniformly along the z-axis segregated into three distinct layers. The midplane acted as an unstable equilibrium, while the top and bottom walls served as stable equilibria that accumulated over 48% of the particles near each boundary (Fig. 2a, 2b). Physical experiments evaluating particle distributions at the device outlet successfully corroborated this phenomenon by confirming the formation of two distinct focal planes. Although numerical simulations of neutrally buoyant particles predicted symmetric equilibrium planes, experimental observations revealed a slightly reduced axial separation distance. This discrepancy was due to the predictable downward gravitational settling of the denser polystyrene particles.

**Figure 2.**
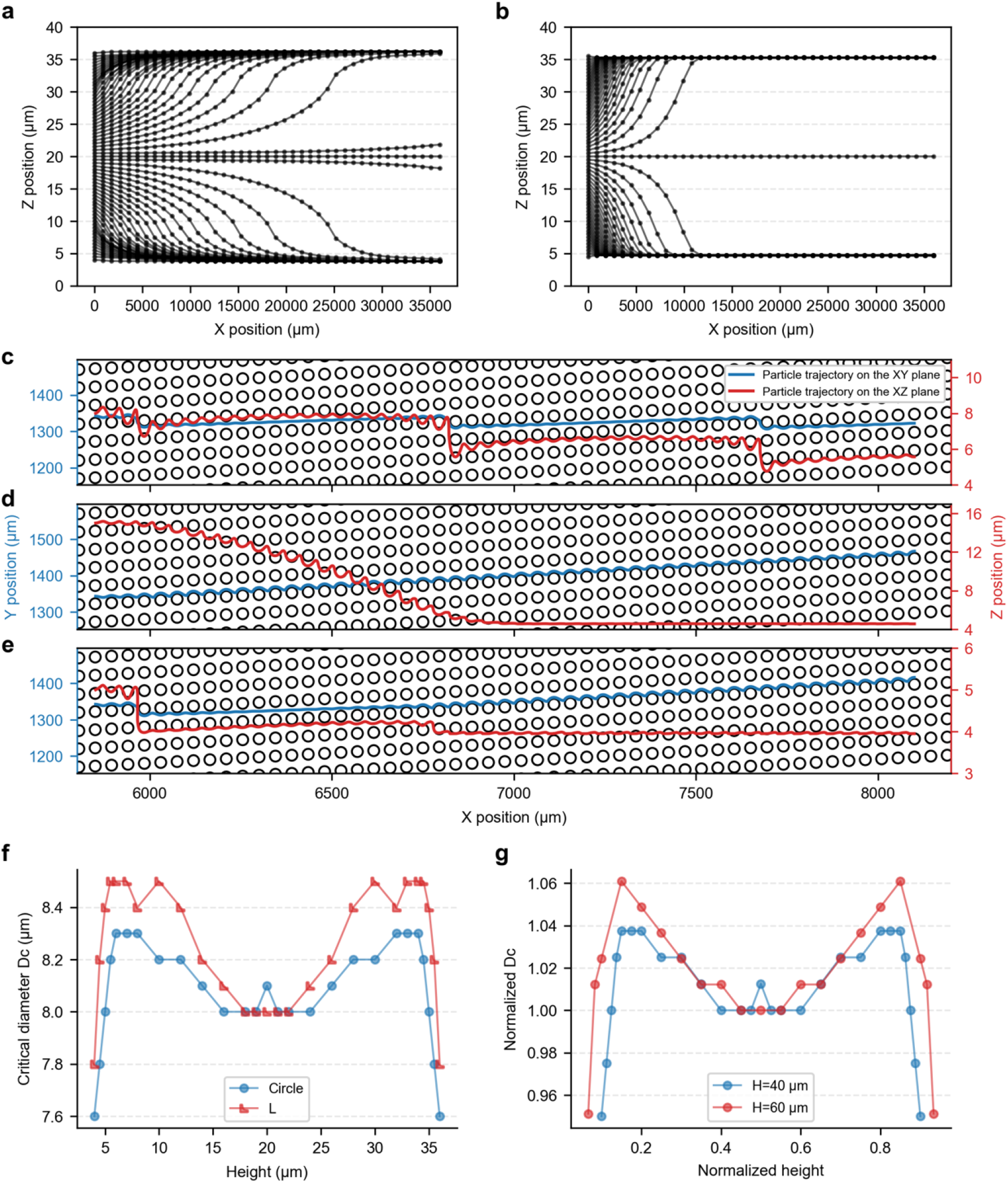
Axial distributions of the critical diameter and 3D particle migration. (**a-b**) Evolution of the axial particle distribution along the flow direction (x-axis) for **a** 7 µm and **d** 9 µm particles. (**c-e**) Projected particle trajectories on the xy-plane and xz-plane for an individual **c** 7 µm particle, **d** 9 µm particle, and **e** 7.8 µm particle. (**f**) Axial distribution of the critical diameter (*D*_*c*_) along the z-axis for DLD devices featuring circular and L-shaped pillars. (**g**) Normalized *D*_*c*_ as a function of the normalized channel height for circular pillar arrays with total height of 40 µm and 60 µm.

Second, extracting the high-resolution axial distribution revealed that the *D*_*c*_ exhibited a distinct M-shaped profile along the z-axis for both circular and L-shaped geometries (Fig. 2f). Referencing the midplane baseline, *D*_*c*_ increased to a local maximum before decaying sharply near the bounding walls, resulting in an overall axial variation of approximately 9%. Increasing the channel height from 40 µm to 60 µm preserved this M-shaped profile while widening the fluctuation amplitude to 11% (Fig. 2g). This axial heterogeneity categorized particle trajectories into three size-dependent regimes. Particles smaller than the minimum *D*_*c*_ strictly followed the zigzag mode, whereas particles larger than the maximum *D*_*c*_ exclusively followed the displacement mode. Meanwhile, intermediate particles exhibited highly position-dependent behavior, with continuous vertical migration driving them across varying local critical diameters. This caused their transport to dynamically alternate between zigzag and displacement trajectories, definitively explaining the macroscopic mixed mode.

Based on these critical discoveries regarding the M-shaped *D*_*c*_ distribution and the accumulation of particles at stable vertical equilibria, we strategically selected coordinates near the upper or lower channel boundaries as the initial starting points for our subsequent 3D FSI simulations. This informed approach ensured that the predictive modeling accurately reflected the true physical distribution of particles within the microfluidic chip.

### 3D FSI simulation achieves superior trajectory prediction

To rigorously validate the proposed simulation framework, we simulated particle trajectories across four distinct DLD geometries including circular, square, drop-shaped, and L-shaped pillars. We employed a rigid-body approximation due to the high stiffness of the particles (Young’s modulus E ≈ 1 − 3 GPa). These specific pillar shapes were selected to introduce varying degrees of flow anisotropy and streamline curvature. We conducted complete inlet-to-outlet fluid simulations of DLD devices composed of a 50 × 1000 pillar array. The FSI framework successfully reproduced the canonical DLD transport modes with high fidelity. These modes comprised two distinct trajectories: a zigzag motion with near-zero net lateral migration, and a displacement motion with significant lateral shift along the array’s inclination angle. Our simulations successfully computed these distinct trajectories for specific particle diameters within the large-scale arrays.

Quantitative comparison confirmed an excellent match between *in silico* predictions and experimental observations. As illustrated in Fig. 3, the simulated trajectories for both displacement-mode and zigzag-mode particles aligned closely with experimental tracks across all four DLD geometries. Notably, the path deviation error remained consistently below 1 µm, representing approximately 5% of the 20 µm gap size. The robustness of the 3D model was particularly evident as it accurately predicted trajectories within L-shaped and drop-shaped arrays, where complex wake structures significantly influenced particle motion. Supplementary Movie 2 further substantiated the precision of the framework by demonstrating how the simulation distinguished trajectory variations originating from distinct initial positions. When benchmarked against high-speed videography, the model faithfully reproduced the fine-scale dynamics of particle deflection, accurately capturing the specific lateral motions required to bypass pillar obstacles within different zigzag-mode tracks. Collectively, these results demonstrated that the fully coupled 3D FSI framework successfully resolved the essential physics of particle transport across a diverse and complex microfluidic design space.

**Figure 3.**
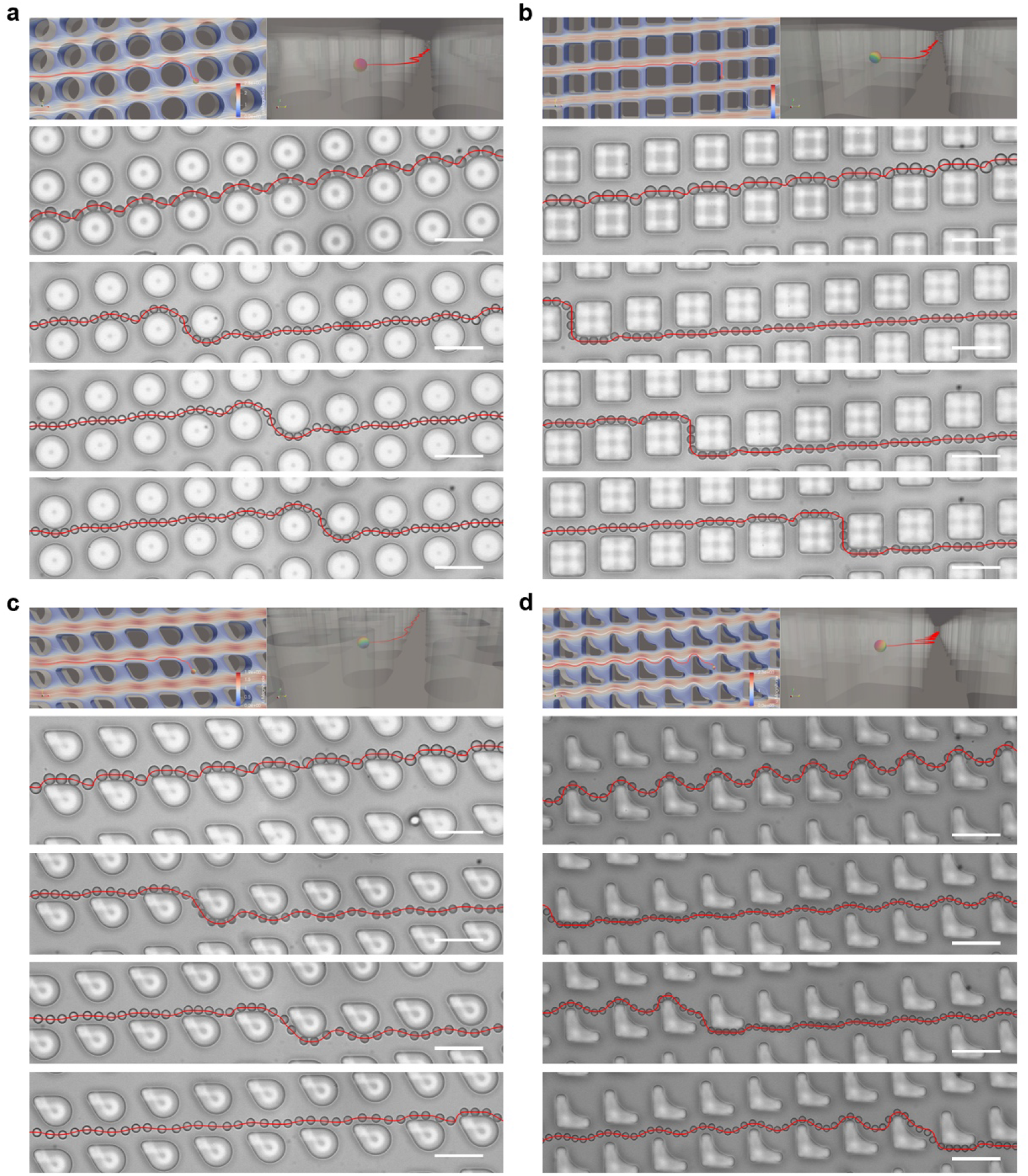
Comparison of experimental and simulated particle trajectories across four distinct DLD pillar geometries. The panels illustrate the sequential positions of particles navigating arrays composed of (**a**) circular, (**b**) square, (**c**) drop-shaped, and (**d**) L-shaped micropillars. In each subfigure, the single trajectory exhibiting the displacement mode corresponds to a large particle (*D*_*p*_ > *D*_*c*_), while the three trajectories exhibiting the zigzag mode correspond to smaller particles (*D*_*p*_ < *D*_*c*_). The excellent alignment between the simulated paths (lines) and experimental tracking (sequential positions) demonstrates the FSI framework’s versatility in handling complex pillar shapes. Scale bars represent 50 µm.

### Prediction and validation of critical diameter

Following the trajectory validation, we employed an algorithm to predict the critical diameter (*D*_*c*_) for each DLD structure. Because the critical diameter represented the defining metric of the device, we determined the predicted *D*_*c*_ by sweeping particle diameters in 0.5 µm increments to identify the transition threshold between the zigzag and displacement modes. To experimentally validate these predictions, we executed a systematic size-sweep protocol by introducing monodisperse microspheres into the microfluidic device in corresponding 0.5 µm increments. Using a dual-inlet sheath flow design, we spatially categorized the outlet positions and compiled the lateral coordinates into distribution histograms to calculate a migration fraction for each particle size. This rigorous statistical approach revealed an experimental transition bandwidth rather than a singular sharp cutoff. Within this interval, the transport behavior shifted from a predominant zigzag mode exhibiting a migration fraction below 20% to a displacement mode exceeding 80%. This specific particle size bandwidth defined the effective experimental critical diameter and accurately captured the mixed-mode phenomenon reported in previous studies^22^. Crucially, our 3D FSI framework accurately predicted this empirical transition bandwidth by explicitly resolving the M-shaped axial fluctuation of local critical diameters and dynamic vertical particle migration.

The 3D FSI framework was utilized to screen 24 distinct DLD structures, comprising eight pillar geometries (circular, drop, L-shaped, inverted triangular, diamond, H-shaped, square, and triangular), each designed with three target critical diameters (*D*_*c*_ = 8, 10, 12 μm). Validation was conducted by comparing experimental measurements against our 3D FSI model and three benchmark models: the empirical formula derived by Davis based on experiments of DLD devices with circular micropillars^39^ (*D*_*c*_ = 1.4Gϵ^0.48^), a 3D computational fluid dynamics (CFD) model relying on massless streamline tracing^40^, and a 2D FSI model utilizing our FSI framework. The 2D FSI model shared identical governing equations and numerical strategies with the 3D framework, differing only by assuming an infinite z-axis depth and treating solids as infinite cylinders.

As illustrated in Fig. 4, the proposed 3D FSI model demonstrates superior accuracy in all cases. The predicted *D*_*c*_ intervals align precisely with experimental measurements across the 24 sample points, yielding a coefficient of determination (R^2^) of 0.98 and a root-mean-square error (RMSE) of 0.38 µm, which is less than 5% of the predicted *D*_*c*_ (Fig. 4d). In contrast, all other models exhibit significant discrepancies. Because the empirical formula originates from circular pillars with isotropic array spacing (equal center-to-center distances in the x and y directions), it yields reasonable agreement for symmetric structures such as square and triangular pillars. However, it deviates significantly when applied to complex geometries, resulting in poor predictive accuracy (R^2^ = 0.14, RMSE = 1.60 µm, Fig. 4a). The 2D FSI method systematically overestimates *D*_*c*_ and yields the largest overall error (R^2^ = 0.09, RMSE = 6.55 µm, Fig. 4c), failing most dramatically in the triangular geometry. This discrepancy arises because the 2D approximation treats particles as infinite cylinders, thereby overestimating the blockage effect compared to 3D reality. Similarly, the 3D fluid-only CFD model treats particles as passive tracers and neglects their influence on the fluid. Consequently, it systematically underestimates the critical diameter, underscoring the necessity of fully resolving coupled fluid-structure interactions (R^2^ = 0.83, RMSE = 3.63 µm, Fig. 4b).

**Figure 4.**
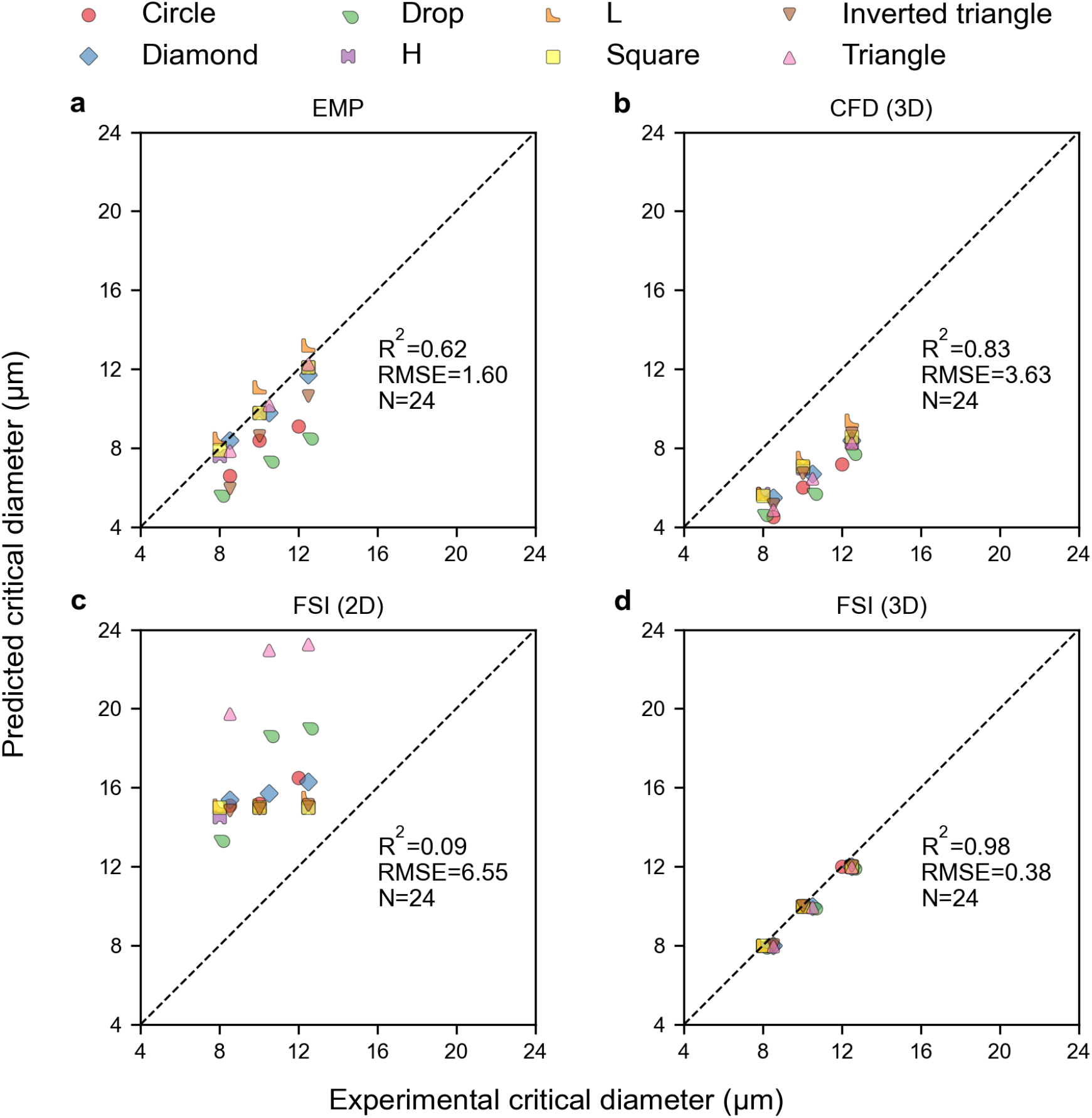
Comparison of predicted versus experimental critical diameters across four different modeling approaches. The plots illustrate the correlation between experimental data and predictions derived from (**a**) the empirical formula derived by Davis^39^ (EMP), (**b**) a 3D computational fluid-only simulation framework^40^ ignoring particle-fluid coupling, (**c**) our proposed 2D FSI simulation framework, and (**d**) our proposed 3D FSI simulation framework. The diagonal dashed line indicates perfect agreement between the predicted and experimental values (y=x). Pillar shapes are represented by marker.

### Application to biological cells

Moving beyond synthetic beads, we explored the framework’s applicability to biological entities by simulating the separation of HT-29 human colorectal adenocarcinoma cells. Although cells were inherently viscoelastic, HT-29 cells exhibited a relatively high stiffness (Young’s modulus E ≈ 0.5 − 1.0kPa) compared to blood cells. Under the hydrodynamic conditions typical of DLD sorting, characterized by low Reynolds numbers and moderate shear, the Capillary Number (*C*_*a*_) for tumor cells remained low (*C*_*a*_ < 0.1). In this regime, fluid shear stress was insufficient to cause significant cellular deformation. Consequently, steric exclusion effects dominated over deformation-induced lift. Based on this physical insight, we modeled the tumor cells as rigid spheres with an effective hydraulic radius.

We validated the 3D FSI framework by comparing simulated rigid particle trajectories against high-speed microscopy of tumor cells (Fig. 5). For cells measuring 9 µm in diameter within circular pillar arrays, the model accurately reproduced the zigzag mode with a trajectory deviation of less than 1 μm (Fig. 5a), maintaining an error of under 2 μm deviation even during complex mid-section rotations (Fig. 5b). In drop-shaped pillar geometries, the framework successfully captured the transport transition, accurately predicting zigzag motion for 7 µm cells (Fig. 5c) and the displacement mode for 12 µm cells (Fig. 5d). Supplementary Movie 3 further reinforced this validation. The video initially projected simulated trajectories for rigid spheres launched from various starting positions. It subsequently displayed the actual experimental tracks of live cells originating from these identical coordinates. Although a localized peak deviation of 4 µm occurred as the 7 µm cell bypassed a pillar, this discrepancy was likely attributable to the simplified rigid-body assumption neglecting cellular deformability under high shear stress near the pillar boundaries. Nevertheless, the framework successfully captured the predominant transport behavior. Crucially, despite minor path variations, the simulation accurately predicted the bifurcation decisions of the cells and correctly identified the migration mode, distinguishing between zigzag and displacement patterns. This high degree of qualitative and modal consistency confirmed the framework’s reliability for predictive DLD design.

**Figure 5.**
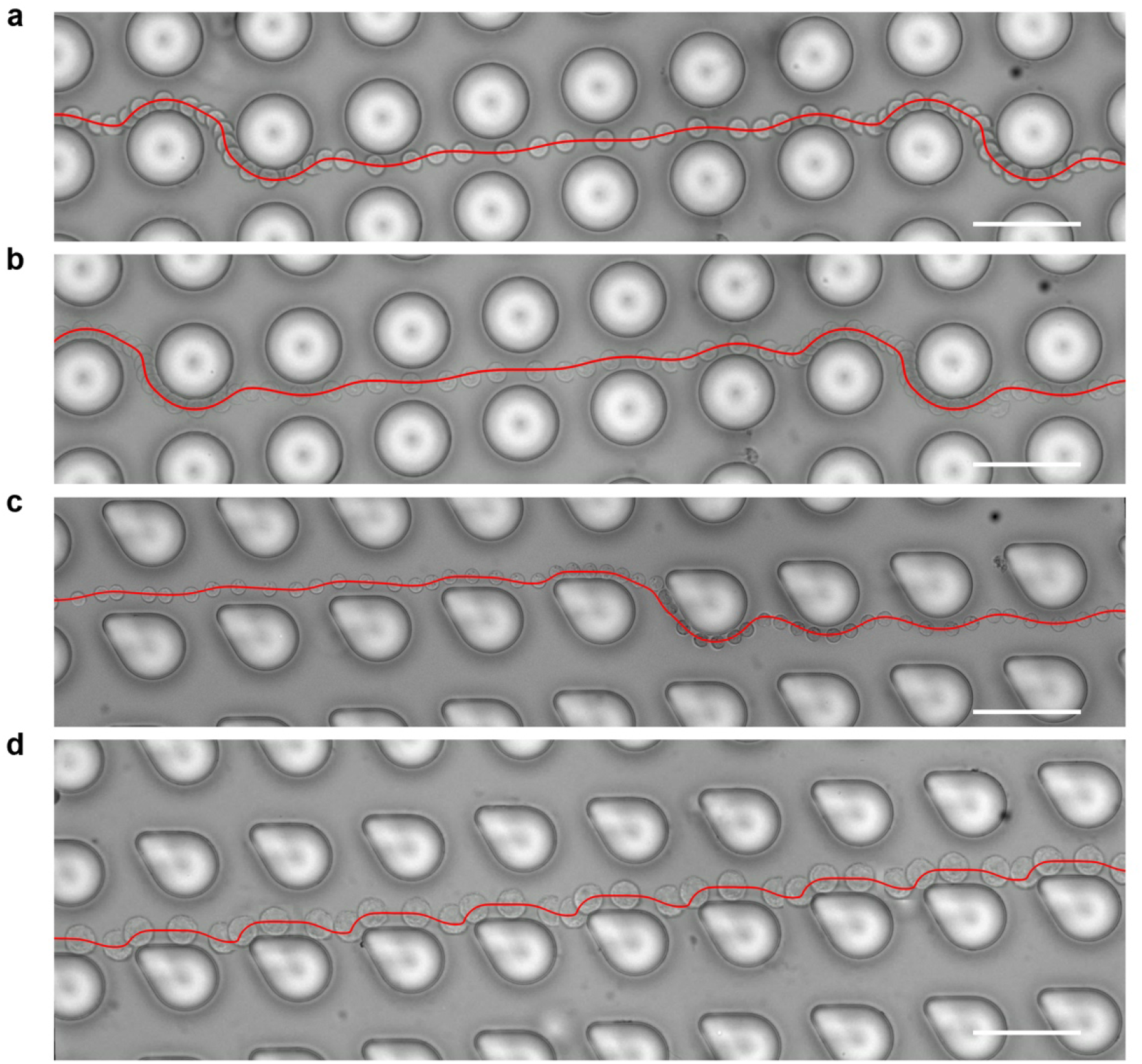
Comparison of experimental tumor cell and simulated particle trajectories in DLD pillar geometries. (**a**) Trajectory of a 9 µm cell in a circular pillar DLD (zigzag mode), with a deviation from the simulated path of < 1 μm. (**b**) Trajectory of a 9 µm cell in a circular pillar DLD (zigzag mode); errors are < 1 μm at the terminal sections and < 2 μm in the middle. (**c**) Trajectory of a 7 µm cell in a drop-shaped pillar DLD (zigzag mode); deviations are < 1 μm initially, reaching a maximum of 4 μm in the mid-section. (**d**) Trajectory of a 12 µm cell in a drop-shaped pillar DLD (displacement mode), with a global error of < 1 μm. Scale bars represent 50 µm.

This result validates the rigid-body engineering approximation for stiff cells, such as circulating tumor cells and epithelial cells, within 3D FSI simulations. It demonstrates that computationally expensive viscoelastic solvers are not always necessary for trajectory prediction if the Capillary number regime is properly identified. With the rigid-body simplification, our framework provides robust and practically relevant guidance for the design and optimization of microfluidic devices intended for cell separation.

### Computational performance and scalability

The 3D FSI framework demonstrated exceptional performance when simulating large-scale, complex architectures. By combining the hierarchical mesh strategy with the localized dynamic mesh technique, we achieved a runtime reduction of two orders of magnitude compared to uniform, globally updated high-resolution meshes (Fig. S3). Furthermore, the synergistic integration of high-order spatial and temporal discretization schemes yielded a transformative leap in computational performance. While the second-order spatial and temporal schemes independently offered 38-fold and 24-fold acceleration, respectively, their combined application resulted in a total speedup of three orders of magnitude compared to a conventional first-order baseline (Fig. S4). Collectively, these algorithmic innovations accelerated the overall computational speed by five orders of magnitude compared to uniform meshes utilizing first-order schemes, thereby rendering full-device simulations highly feasible. For instance, simulating a comprehensive particle trajectory within a specific deterministic lateral displacement candidate geometry can now be completed in just 30 hours. Because these individual geometric evaluations can be routinely executed in parallel, this ultra-high efficiency establishes a robust foundation for high-throughput automated design optimization.

### High-throughput screening and predictive structural optimization

Leveraging this unprecedented computational efficiency, the framework transcends retrospective validation to function as a powerful instrument for predictive structural optimization and rapid design-space exploration. To systematically optimize separation arrays for a specific target critical diameter, researchers can employ a grid search algorithm to evaluate diverse parameter combinations. These iterations can encompass simultaneous variations in pillar geometry, gap size, row-shift fraction, row-shift angle, and device depth. As a demonstration of high-throughput screening, the execution of a comprehensive batch of 48 distinct configurations required 30 hours utilizing 8 cloud servers (each equipped with dual Intel Xeon 8260M CPUs and 512GB RAM). This workflow represented a computational investment of 11,520 CPU hours at a minimal cost of 322 RMB (based on a rate of 0.028 RMB per CPU-hour). Crucially, this cloud computing architecture inherently supports massive parallel testing, empowering engineers to increase the scale of evaluated designs without extending the overall completion time. Because the predicted critical diameters exhibit exceptional concordance with experimental measurements, designers can confidently identify the optimal architecture to meet specific separation criteria. Consequently, this highly efficient predictive workflow provides a cost-effective alternative to traditional empirical prototyping, accelerating the development of practical microfluidic systems by enabling rigorous, simulation-driven design iterations.

## Discussion

The transition from empirical trial-and-error prototyping to simulation-driven engineering represents a critical evolutionary step for the maturation of modern microfluidics. Although microfluidic technologies possess transformative potential, their commercial scalability remains severely constrained by a persistent reliance on empirical physical iterations. This study directly addresses this systemic bottleneck by establishing a unified 3D FSI framework, utilizing DLD architectures as a stringent paradigm to validate its practical feasibility. By successfully reconciling the conflict between strict physical fidelity and computational tractability, this methodology fundamentally alters the economics of microfluidic engineering. Rather than functioning merely as a retrospective analytical tool, this highly accelerated infrastructure liberates researchers from traditional hardware limitations. Coupling this robust computational framework with cloud-based parallel computing environments enables the systematic exploration of massive design parameter spaces. This exceptional scalability ensures that evaluating extensive geometric variations simultaneously no longer imposes prohibitive temporal or financial burdens. Because the resulting simulations exhibit extraordinary concordance with physical measurements, designers can confidently optimize complex multi-stage architectures entirely within a virtual environment. The capacity to accurately identify optimal structural configurations before manufacturing a single prototype establishes this framework as a powerful instrument for high-throughput screening and predictive structural optimization, fundamentally overcoming traditional hardware development bottlenecks.

To achieve such robust predictive capabilities, A central finding of this study is that accurately predicting separation dynamics in DLD arrays requires a fully coupled 3D approach. Fluid-only models systematically underestimate the critical diameter because they treat particles as passive tracers and entirely neglect the hydrodynamic blockage effect^40^. In physical devices, the volume of a moving particle perturbs the local pressure field and interacts strongly with the no-slip boundaries at the channel floor and ceiling. This dynamic interaction generates significant wall-induced lift forces that repel particles into higher-velocity central streamlines, thereby expanding their effective hydraulic footprint and altering the bifurcation conditions at the pillar interface. Beyond the necessity of fluid-structure coupling, our results definitively expose the severe limitations of 2D spatial approximations. Because planar models treat particles as infinite cylinders and artificially enforce zero vertical velocity, they inherently fail to capture vertical shear gradients and 3D rotational dynamics. The critical importance of resolving these spatial effects is underscored by the superior predictive accuracy of our framework within anisotropic L-shaped and drop-shaped arrays, where complex secondary flows exist such as Dean vortices. Ultimately, particle separation is governed by an intricate 3D interplay between the physical object and its surrounding hydrodynamic environment. By explicitly resolving these mechanisms, our 3D FSI framework provides an exceptional tool to deeply investigate the fundamental physics of microfluidic separation.

Building upon this foundation, this advanced simulation capability is particularly crucial for elucidating the mechanistic origins of the mixed mode, a phenomenon that has long remained highly debated within the academic community. In both our experiments and the broader literature, particles of specific sizes frequently exhibit a mixed mode, alternating between zigzag and displacement trajectories. Previous 2D-based investigations attributed this behavior to flow lane asymmetry or anisotropic permeability^22,41^, while a recent 3D study proposed a U-shaped *D*_*c*_ distribution along the z-axis^42^. Our fully coupled 3D FSI model reveals that the true axial *D*_*c*_ distribution is distinctly M-shaped as depicted in Fig. 2a. Coupled with the discovery of dynamic longitudinal migration, this physical insight definitively resolves the genesis of the mixed mode. For intermediate-sized particles falling within this M-shaped fluctuation range, their instantaneous separation mode becomes fundamentally tied to their z-coordinate. As these particles migrate axially through the microchannel, they actively traverse spatial regions characterized by varying local *D*_*c*_. Consequently, moving into a region where the local *D*_*c*_ exceeds the particle size triggers zigzag transport, whereas entering a zone with a lower local *D*_*c*_ forces displacement. This dynamic and axially driven threshold-crossing process manifests macroscopically as the mixed separation mode. Furthermore, practical experimental conditions such as inherent particle polydispersity, fabrication tolerances, and pressure-induced device deformation inevitably compound this effect by locally perturbing both the flow fields and the critical thresholds. Our 3D FSI simulation provides a deterministic explanation for previously ambiguous mixed-mode sorting behaviors and advances the understanding of separation physics from a three-dimensional perspective. Ultimately, only a predictive digital design methodology driven by fully coupled 3D FSI simulations can explicitly harness these complex hydrodynamic effects to design microfluidic chips with unprecedented separation precision.

Translating this unprecedented precision from synthetic particles to complex biological entities, our framework was applied to tumor cells, providing validation for rigid body engineering approximations under specific biological conditions. Although cells are inherently viscoelastic, our findings suggest that full multiphysics modeling of deformability is not a prerequisite for accurate trajectory prediction when specific physical conditions are met. For instance, the HT-29 tumor cells tested here are characterized by a relatively high stiffness modulus (E ≈ 0.5 − 1.0kPa) and operate within a low Capillary number regime (*C*_*a*_ < 0.1) in our DLD devices. Under these conditions, hydrodynamic shear stresses are insufficient to induce significant cellular deformation, thereby allowing steric exclusion to dominate over deformation-induced lift forces. The excellent agreement observed between our rigid-body simulations and experimental tumor cell trajectories (Fig. 5 and Movie S3) supports a practical hierarchical design strategy where computationally expensive deformable cell models are reserved for high-Capillary number regimes or highly flexible cells like erythrocytes. Conversely, rigid-body FSI serves as a rapid and accurate alternative for stiffer cell lines such as circulating tumor cells, significantly accelerating the design cycle for oncological sorting applications without compromising predictive fidelity.

While accurately modeling these localized cell-pillar interactions is vital, realizing true digital automation also necessitates scaling these simulations to encompass entire devices. Standard simulation methods have historically relied on periodic boundary conditions to simulate a single unit cell, thereby implicitly assuming an infinite and uniform lattice. While mathematically convenient, this approach fundamentally fails to capture boundary effects that dictate real-world chip performance across critical non-periodic regions such as inlets, outlets, and sidewalls. Our computational framework overcomes this limitation by leveraging a hierarchical mesh strategy to model centimeter-scale devices entirely without assuming structural periodicity. By employing a coarse-mesh macroscopic simulation to establish accurate boundary conditions for the high-resolution analysis of any specific region of interest, the framework successfully resolves arbitrary local domains. As demonstrated by the simulation of the device periphery in Fig. S5, the flow field near the physical edge diverged significantly from the central region and exhibited a marked increase in fluid velocity. Crucially, the model accurately captured particle trajectories within these boundary-adjacent arrays and demonstrated excellent agreement with experimental tracks recorded via high-speed imaging. Because these critical boundary dynamics are inherently missed by conventional unit-cell models, accurately resolving such non-periodic regions provides a profound engineering advantage. Our framework facilitates the rigorous optimization of peripheral edges and inter-stage transition zones to ensure that particles are correctly equilibrated before entering downstream separation arrays. This unique capability to model arbitrary chip locations advances the field from isolated component optimization to integrated system engineering, establishing a reliable computational foundation for robust diagnostic platforms.

Executing such expansive system-level models naturally demands an unprecedented leap in algorithmic efficiency. Historically, the prohibitive computational cost of 3D FSI has restricted its use to supercomputing environments. We have overcome this barrier by synergizing three architectural innovations that make high-fidelity simulation accessible on standard cloud servers. First, our hierarchical mesh strategy efficiently decouples the device scale from the particle scale, concentrating high resolution exclusively in physically critical regions. Second, the adoption of high-order spatio-temporal algorithms significantly mitigates numerical diffusion, permitting the use of larger time steps without sacrificing fidelity. Third, a localized dynamic mesh-updating strategy avoids the prohibitive costs of global remeshing while maintaining precise interface tracking. Collectively, these innovations reduce runtime by five orders of magnitude compared to uniform meshes utilizing first-order schemes. This efficiency enables full trajectory simulations to be completed in days, effectively transforming 3D FSI into a prospective instrument for engineering design.

Although this framework signifies a major advancement in microfluidic simulation, its current reliance on a rigid-body approximation restricts its applicability to stiffer cells. To accurately model highly deformable entities like red blood cells and extracellular vesicles, future iterations will integrate viscoelastic cell models to explicitly resolve the coupled dynamics of fluid flow, solid translation, and structural deformation. In parallel, ongoing algorithmic developments aim to combine direct matrix solvers such as MUMPS with iterative multigrid methods. This integration will accelerate computations by an additional order of magnitude to further expand high-throughput design capabilities. Nevertheless, the fully coupled 3D framework robustly captures the fundamental hydrodynamics of particle transport across arbitrarily complex geometries, making it exceptionally suitable for a wide range of passive label-free separation technologies. Moving beyond deterministic lateral displacement, future deployments will utilize this framework to deeply investigate hidden physical mechanisms and perform predictive structural optimization for advanced devices governed by the principles of inertial microfluidics, pinched flow fractionation, and viscoelastic microfluidics.

In conclusion, this work delivers the first experimentally validated and computationally efficient 3D FSI framework capable of resolving the multi-scale physics governing microfluidic separation. By demonstrating that predictive digital design can match the fidelity of physical experiments, we provide a transformative toolset for the microfluidics community. Importantly, the framework provides profound insights into the hidden 3D hydrodynamic mechanisms underlying deterministic lateral displacement systems, thereby facilitating the development of exceptional precise separation strategies. By providing a robust computational infrastructure for virtual prototyping, the framework enables a paradigm shift from slow empirical prototyping to rapid *in silico* optimization. Ultimately, by facilitating the precise digital replication of complex fluid-structure phenomena, this technology empowers the transition toward simulation-driven engineering and accelerates the commercial deployment of next-generation lab-on-a-chip innovations.

## Methods

### Computational workflow for multi-scale fluid-structure interaction

Simulating particle dynamics within microfluidic arrays involves significant scale disparity. To avoid the prohibitive computational costs of continuous global remeshing, we implemented a localized dynamic mesh-updating algorithm. This approach effectively decoupled the macroscopic channel hydrodynamics from the transient fluid-structure interaction (FSI) dynamics. We executed the computational workflow in three coupled phases.

### Background flow pre-computation

To establish the macroscopic hydrodynamic environment, a full-scale 3D tetrahedral mesh encompassing the entire DLD array (50 rows × 1000 columns) was generated. This global background mesh (Ω_bg_) utilized a hierarchical resolution strategy: coarse elements were deployed in the far-field, medium refinement was applied within anticipated particle trajectory zones, and second-order elements were imposed at all pillar boundaries to accurately capture shear gradients (Figure 1b). We solved the steady-state incompressible Navier-Stokes equations on this static domain utilizing a P2-P1 (Taylor-Hood) finite element method:

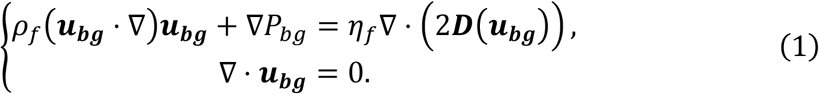

Here,*ρ*_***f***_ stands for the fluid density, *η*_*f*_ is the viscosity of the fluid, and 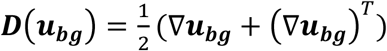 represents the fluid strain-rate tensor. The boundary conditions comprised a uniform velocity profile at the channel inlet, a do-nothing boundary condition at the outlet, and no-slip conditions on all other walls. This global solution yielded the background velocity (***u***_***bg***_) and pressure fields (*P*_*bg*_), which were utilized for subsequent localized FSI simulation.

### Localized FSI domain construction and monolithic time integration

To resolve the transient fluid-solid interfacial dynamics, a high-resolution, localized computational sub-domain (Ω_*loc*_) spanning a 4 × 4 pillar neighborhood was dynamically constructed around the target particle. This localized domain featured a body-fitted matching tetrahedral mesh at the fluid-solid interface, with dynamic adaptive refinement strictly confined to the particle vicinity and second-order elements applied to all boundaries (Figure 1d). At each time increment, the pre-computed macroscopic velocity field was spatially interpolated onto the exterior boundaries (∂Ω_*loc*_) to enforce transient Dirichlet boundary conditions:

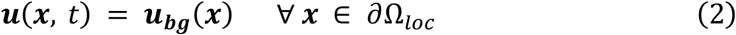

Within this localized domain, the fluid-structure system was modeled as the coupling between an incompressible Newtonian fluid and an incompressible hyperelastic solid. The fluid motion was governed by the Navier–Stokes equations:

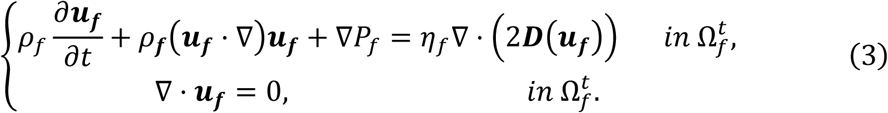

The solid was described using a neo-Hookean constitutive model formulated in terms of the left Cauchy–Green deformation tensor ***B***:

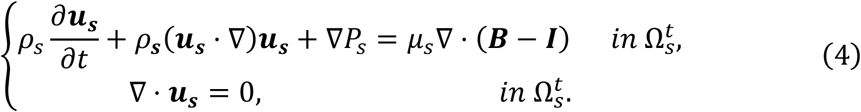

where 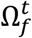 and 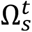 denote the fluid and solid domains, ***u***_***f***_ and ***u***_***s***_ are the velocities of the fluid and solid, *P*_*f*_ and *P*_*s*_ represent the corresponding pressures, *μ*_*s*_ is the shear modulus of the solid, and ***I*** is the identity matrix. The two phases occupied complementary time-dependent subdomains Ω_*loc*_ and were strongly coupled through the kinematic continuity of velocity and the dynamic equilibrium of stresses at the fluid-solid interface Γ^*t*^:

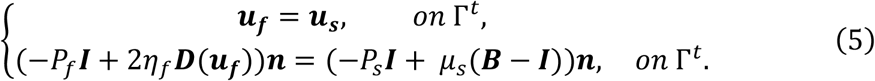

Here, ***n*** represents the unit normal vector of the interface Γ^*t*^. The evolution of the structural deformation was tracked via a transport equation for the deformation tensor, ensuring consistency with the underlying kinematics:

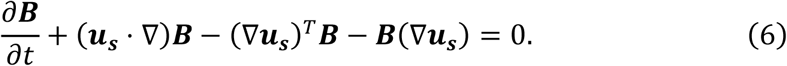

This sharp-interface formulation enabled accurate resolution of interfacial stresses and deformation while preserving a unified monolithic treatment of the coupled system.

Within Ω_*loc*_, an Arbitrary Lagrangian–Eulerian (ALE) formulation^43^ was employed to seamlessly track the moving fluid-solid interface and manage the corresponding mesh updates. The mesh velocity ***w***_***t***_ was computed in a decoupled manner by solving a pseudo-elastic extension problem subject to Dirichlet boundary conditions prescribed by the structural velocity at the fluid-solid interface and homogeneous constraints at the outer boundary. To prevent volumetric distortion, ***w***_***t***_ was also constrained to satisfy a divergence-free condition where ∇ ⋅ *w*_*t*_ = 0.

Incorporating the mesh velocity into the governing equations yielded the total continuous system:

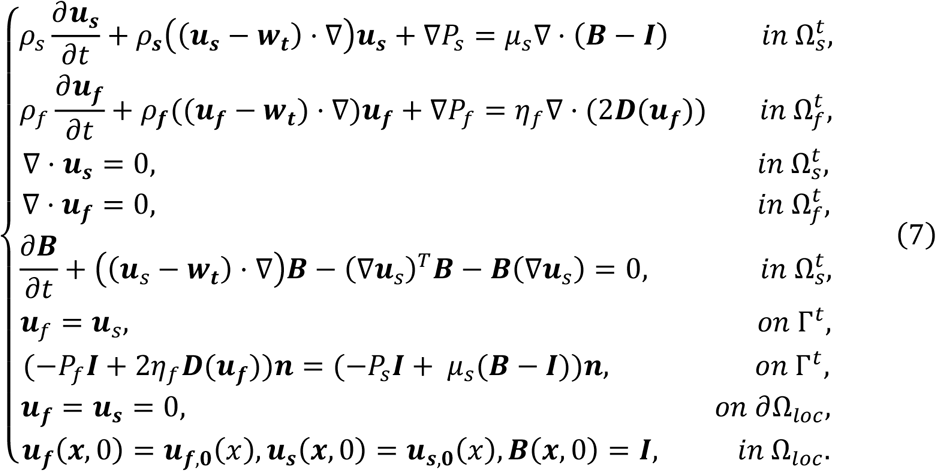

Following the implicit convergence of the monolithic fluid-structure system at each time step, the spatial trajectory of the solid particle was then explicitly advanced using a second-order Runge-Kutta (RK2) scheme based on the computed solid kinematics. Technical details regarding the discretization algorithms and the implementation are provided in our recent work^44^.

### Dynamic adaptive remeshing and state projection

As the particle underwent continuous displacement, the surrounding ALE mesh inevitably deformed. To prevent severe element distortion, an adaptive remeshing protocol continuously monitored local mesh quality metrics. Once a predefined distortion threshold was exceeded, a new optimized local mesh (∂Ω^′^ _*loc*_) was generated at the updated particle coordinates. Transient physical fields were then projected from the distorted mesh onto the new grid. In spatially overlapping regions, state variables were transferred via high-order interpolation to preserve interfacial fidelity. For non-overlapping regions entering the sub-domain, the fluid velocity was interpolated from *u*_*bg*_. This seamless dynamic mapping sustained high-order accuracy at the fluid-solid interface while enabling continuous trajectory tracking across the ultra-long microfluidic channel.

### Microfluidic Device Fabrication

The DLD devices were fabricated using standard soft lithography techniques. Master molds were created by spin coating a layer of negative photoresist (SU-8 3050, MicroChem) on 4-inch silicon wafers and subsequently soft-baked at 65°C and 95°C. Pillar array patterns encompassing circular, drop-shaped, L-shaped, inverted triangular, diamond, H-shaped, square, and triangular geometries were transferred to the wafer via UV exposure using a mask aligner (Karl Suss MA6) followed by development. To create the microfluidic devices, Polydimethylsiloxane (PDMS) prepolymer (RTV 615, GE) was mixed with a curing agent at a 10:1 (w/w) ratio and degassed in a vacuum chamber for 30 minutes. The mixture was cast over the silanized SU-8 master and cured at 80°C for 45 minutes. Upon completion of curing, the PDMS replicas were peeled off and punched with a biopsy punch to form inlet and outlet ports. Final assembly was achieved by sealing the device against a clean glass slide treated with oxygen plasma to ensure irreversible bonding.

#### Rigid Particle Preparation

Monodisperse fluorescent polystyrene microspheres (Suzhou Nanomicro Technology Co., Ltd.) with diameters ranging from 7 µm to 15 µm were used for trajectory validation and critical diameter determination. The beads were suspended in an aqueous buffer containing 0.1% Tween-20 to prevent aggregation and adhesion to channel walls.

#### Cell Culture

Human colorectal adenocarcinoma cells (HT-29) were obtained from the American Type Culture Collection (ATCC). Cells were cultured in Dulbecco’s Modified Eagle Medium (DMEM) supplemented with 10% Fetal Bovine Serum (FBS) and 1% penicillin-streptomycin in a humidified incubator at 37°C with 5% CO_2_. Prior to experiments, cells were harvested using 0.25% Trypsin-EDTA and resuspended in Phosphate Buffered Saline (PBS). The stiffness of HT-29 cells allows for a rigid-body approximation under the low shear stress conditions typical of the tested DLD arrays.

#### Experimental Setup and Image Acquisition

To quantify particle dynamics, the microfluidic device was mounted on the stage of an inverted fluorescence microscope (Nikon Eclipse Ti2) equipped with a high-speed sCMOS camera (Kinetix, Teledyne Photometrics). Fluid flow was driven by a high-precision pressure controller (Elveflow OB1 MK4) to maintain stable, pulse-free flow rates. For trajectory analysis, image sequences were captured at frame rates of approximately 900 fps, with the particle velocity of approximately 10 mm/s.

#### Trajectory Tracking

Experimental videos were processed using noise reduction and spatial segmentation to isolate individual targets. A particle tracking velocimetry algorithm then reconstructed the motion trajectories by linking particle centroids across consecutive frames. This computational approach enabled the precise extraction of lateral coordinates as particles traversed the observation window.

#### Measurement of Critical Diameter

The critical diameter was determined experimentally through a systematic size-sweep protocol by introducing monodisperse polystyrene microspheres in 0.5 µm diameter increments. Utilizing a dual-inlet sheath flow architecture, we divided the outlet into a non-displaced region for the initial sheath flow and a displaced region for the upper buffer. The lateral outlet coordinates of all tracked particles were compiled to generate comprehensive spatial distribution histograms. We then evaluated the overall separation efficiency by defining a migration fraction *ϕ*, which represents the percentage of particles successfully reaching the displaced region relative to the total number of tracked observations.

#### Statistical Analysis and Mode Classification

Statistical thresholds based on the calculated migration fraction *ϕ* were applied to robustly classify the transport behaviors. A migration fraction *ϕ* < 20% statistically indicated a predominant zigzag mode, confirming that the particle size is smaller than the critical diameter. Conversely, a fraction *ϕ* > 80% denoted a displacement mode, demonstrating that the particle size surpasses the critical threshold. Intermediate values where 20% < *ϕ* < 80% defined the separation transition bandwidth. Consequently, we identified the effective critical diameter as the specific particle size interval capturing this functional transition from the zigzag to the displacement mode. All experiments were performed in triplicate to ensure reproducibility. Finally, we assessed the predictive accuracy of the simulation framework by calculating the root-mean-square error between simulated and experimental critical diameters.

## Acknowledgements

This work is supported by the funding of Shenzhen Raymind Biotechnology Co., Ltd.

## Author contributions

J. Han and Y. Chen conceived the research and designed the study. L. Shen led the model development, with algorithmic implementation contributed by Y. Zhang, Y. X. Chen, and P. Wen. C. Wang, P. Sun, S. Gong, and J. Xu contributed to technical discussions and refinement of the algorithms. X. Ding and Y. Zhang designed the experiments and analyzed the results. Y. Chen, L. Shen, Y. Zhang, and J. Han wrote and edited the manuscript. All authors reviewed and approved the manuscript prior to submission.

## Conflict of interest statement

The authors declare no competing interest.

**Supplementary information** is available for this paper at the attachments.

## Reference

1. Huang, L. R., Cox, E. C., Austin, R. H. & Sturm, J. C. Continuous Particle Separation Through Deterministic Lateral Displacement. Science 304, 987–990 (2004).

2. Salafi, T., Zhang, Y. & Zhang, Y. A Review on Deterministic Lateral Displacement for Particle Separation and Detection. Nano-Micro Letters 11, 77 (2019).

3. Beech, J. P., Holm, S. H., Adolfsson, K. & Tegenfeldt, J. O. Sorting cells by size, shape and deformability. Lab Chip 12, 1048–1051 (2012).

4. Xavier, M. et al. Label-free enrichment of primary human skeletal progenitor cells using deterministic lateral displacement. Lab Chip 19, 513–523 (2019).

5. Holm, S. H., Beech, J. P., Barrett, M. P. & Tegenfeldt, J. O. Simplifying microfluidic separation devices towards field-detection of blood parasites. Anal. Methods 8, 3291–3300 (2016).

6. Davis, J. A. et al. Deterministic hydrodynamics: Taking blood apart. Proceedings of the National Academy of Sciences 103, 14779–14784 (2006).

7. Loutherback, K. et al. Deterministic separation of cancer cells from blood at 10 mL/min. AIP Advances 2, 042107 (2012).

8. Ranjan, S., Zeming, K. K., Jureen, R., Fisher, D. & Zhang, Y. DLD pillar shape design for efficient separation of spherical and non-spherical bioparticles. Lab Chip 14, 4250–4262 (2014).

9. Inglis, D. W., Herman, N. & Vesey, G. Highly accurate deterministic lateral displacement device and its application to purification of fungal spores. Biomicrofluidics 4, 024109 (2010).

10. Wunsch, B. H. et al. Nanoscale lateral displacement arrays for the separation of exosomes and colloids down to 20 nm. Nature Nanotechnology 11, 936–940 (2016).

11. Wunsch, B. H. et al. Gel-on-a-chip: continuous, velocity-dependent DNA separation using nanoscale lateral displacement. Lab Chip 19, 1567–1578 (2019).

12. Chen, Y. et al. Concentrating Genomic Length DNA in a Microfabricated Array. Phys. Rev. Lett. 114, 198303 (2015).

13. Hochstetter, A. et al. Deterministic Lateral Displacement: Challenges and Perspectives. ACS Nano 14, 10784–10795 (2020).

14. P. Español & P. Warren. Statistical Mechanics of Dissipative Particle Dynamics. Europhysics Letters 30, 191 (1995).

15. P. J. Hoogerbrugge & J. M. V. A. Koelman. Simulating Microscopic Hydrodynamic Phenomena with Dissipative Particle Dynamics. Europhysics Letters 19, 155 (1992).

16. Zhang, Z., Henry, E., Gompper, G. & Fedosov, D. A. Behavior of rigid and deformable particles in deterministic lateral displacement devices with different post shapes. The Journal of Chemical Physics 143, 243145 (2015).

17. Holm, S. H. et al. Microfluidic Particle Sorting in Concentrated Erythrocyte Suspensions. Phys. Rev. Appl. 12, 014051 (2019).

18. Liu, S. et al. Unraveling the motion and deformation characteristics of red blood cells in a deterministic lateral displacement device. Computers in Biology and Medicine 168, 107712 (2024).

19. Aidun, C. K. & Clausen, J. R. Lattice-Boltzmann Method for Complex Flows. Annual Review of Fluid Mechanics vol. 42 439–472 (2010).

20. Succi, S. The Lattice Boltzmann Equation for Fluid Dynamics and Beyond. (Oxford University Press, 2001). doi:10.1093/oso/9780198503989.001.0001.

21. Vernekar, R. & Krüger, T. Breakdown of deterministic lateral displacement efficiency for non-dilute suspensions: A numerical study. Medical Engineering & Physics 37, 845–854 (2015).

22. Kulrattanarak, T., van der Sman, R. G. M., Schroën, C.G.P.H. & Boom, R. M. Analysis of mixed motion in deterministic ratchets via experiment and particle simulation. Microfluidics and Nanofluidics 10, 843–853 (2011).

23. Reinecke, S. R. et al. DEM-LBM simulation of multidimensional fractionation by size and density through deterministic lateral displacement at various Reynolds numbers. Powder Technology 385, 418–433 (2021).

24. Peskin, C. S. Flow patterns around heart valves: A numerical method. Journal of Computational Physics 10, 252–271 (1972).

25. Yu, Z., Yang, Y. & Lin, J. Lubrication Force Saturation Matters for the Critical Separation Size of the Non-Colloidal Spherical Particle in the Deterministic Lateral Displacement Device. Applied Sciences 12, 2733 (2022).

26. Glowinski, R., Pan, T.-W., Hesla, T. I. & Joseph, D. D. A distributed Lagrange multiplier/fictitious domain method for particulate flows. International Journal of Multiphase Flow 25, 755–794 (1999).

27. Krüger, T., Holmes, D. & Coveney, P. V. Deformability-based red blood cell separation in deterministic lateral displacement devices—A simulation study. Biomicrofluidics 8, 054114 (2014).

28. Wullenweber, M. S., Kottmeier, J., Kampen, I., Dietzel, A. & Kwade, A. Simulative Investigation of Different DLD Microsystem Designs with Increased Reynolds Numbers Using a Two-Way Coupled IBM-CFD/6-DOF Approach. Processes 10, (2022).

29. Wullenweber, M. S., Kottmeier, J., Kampen, I., Dietzel, A. & Kwade, A. Numerical Study on High Throughput and High Solid Particle Separation in Deterministic Lateral Displacement Microarrays. Processes 11, 2438 (2023).

30. Hirt, C. W., Amsden, A. A. & Cook, J. L. An arbitrary Lagrangian-Eulerian computing method for all flow speeds. Journal of Computational Physics 14, 227– 253 (1974).

31. Donea, J., Giuliani, S. & Halleux, J. P. An arbitrary lagrangian-eulerian finite element method for transient dynamic fluid-structure interactions. Computer Methods in Applied Mechanics and Engineering 33, 689–723 (1982).

32. D’Avino, G. Non-Newtonian deterministic lateral displacement separator: theory and simulations. Rheologica Acta 52, 221–236 (2013).

33. Khodaee, F., Movahed, S., Fatouraee, N. & Daneshmand, F. Numerical Simulation of Separation of Circulating Tumor Cells from Blood Stream in Deterministic Lateral Displacement (DLD) Microfluidic Channel. Journal of Mechanics 32, 463–471 (2016).

34. Rezaei, B., Moghimi Zand, M. & Javidi, R. Numerical simulation of critical particle size in asymmetrical deterministic lateral displacement. Journal of Chromatography A 1649, 462216 (2021).

35. Chen, X., Feng, Q. & Zhang, Y. Insights into a deterministic lateral displacement sorting chip with new cross-section micropillars. Chaos, Solitons & Fractals 156, 111884 (2022).

36. Wang, C. et al. A unified-field monolithic fictitious domain-finite element method for fluid-structure-contact interactions and applications to deterministic lateral displacement problems. Journal of Computational Physics 510, 113083 (2024).

37. Liang, Y., Wang, C., Sun, P., Chen, Y. & Han, J. An interface-fitted/fictitious domain-finite element method for the simulation of particles falling in viscous fluid with contact. Computers & Fluids 299, 106685 (2025).

38. Wang, C., Liang, Y., Sun, P., Chen, Y. & Han, J. An energy-stable monolithic interface-fitted/fictitious domain-finite element method for interaction problems of fluid and rigid body with large displacements. Journal of Computational Physics 553, 114720 (2026).

39. Davis, J. A. Microfluidic Separation of Blood Components through Deterministic Lateral Displacement. in (2008).

40. Inglis, D. W., Davis, J. A., Austin, R. H. & Sturm, J. C. Critical particle size for fractionation by deterministic lateral displacement. Lab Chip 6, 655–658 (2006).

41. Kim, S.-C. et al. Broken flow symmetry explains the dynamics of small particles in deterministic lateral displacement arrays. Proceedings of the National Academy of Sciences 114, E5034–E5041 (2017).

42. Chen, J., Huang, X., Xuan, W. & Sun, L. A 3D modeling framework for accurate trajectory-based prediction of critical diameter in deterministic lateral displacement microfluidics. Microsystems & Nanoengineering 12, 78 (2026).

43. Hao, W., Sun, P., Xu, J. & Zhang, L. Multiscale and monolithic arbitrary Lagrangian–Eulerian finite element method for a hemodynamic fluid-structure interaction problem involving aneurysms. Journal of Computational Physics 433, 110181 (2021).

44. Shen, L. et al. A monolithic localized high-order ALE finite element method for multi-scale fluid-structure interaction problems. Preprint at 10.48550/arXiv.2602.02003 (2026).

