## Supplementary Information for "A High-Fidelity 3D Fluid-Structure Interaction Framework for Predictive Microfluidic Design"

This PDF file includes

Supplementary information S1 to S4

Figure S1 to S5

Table S1

Movie S1 to S3

### S1. Microfluidic chip for 3D FSI framework validation

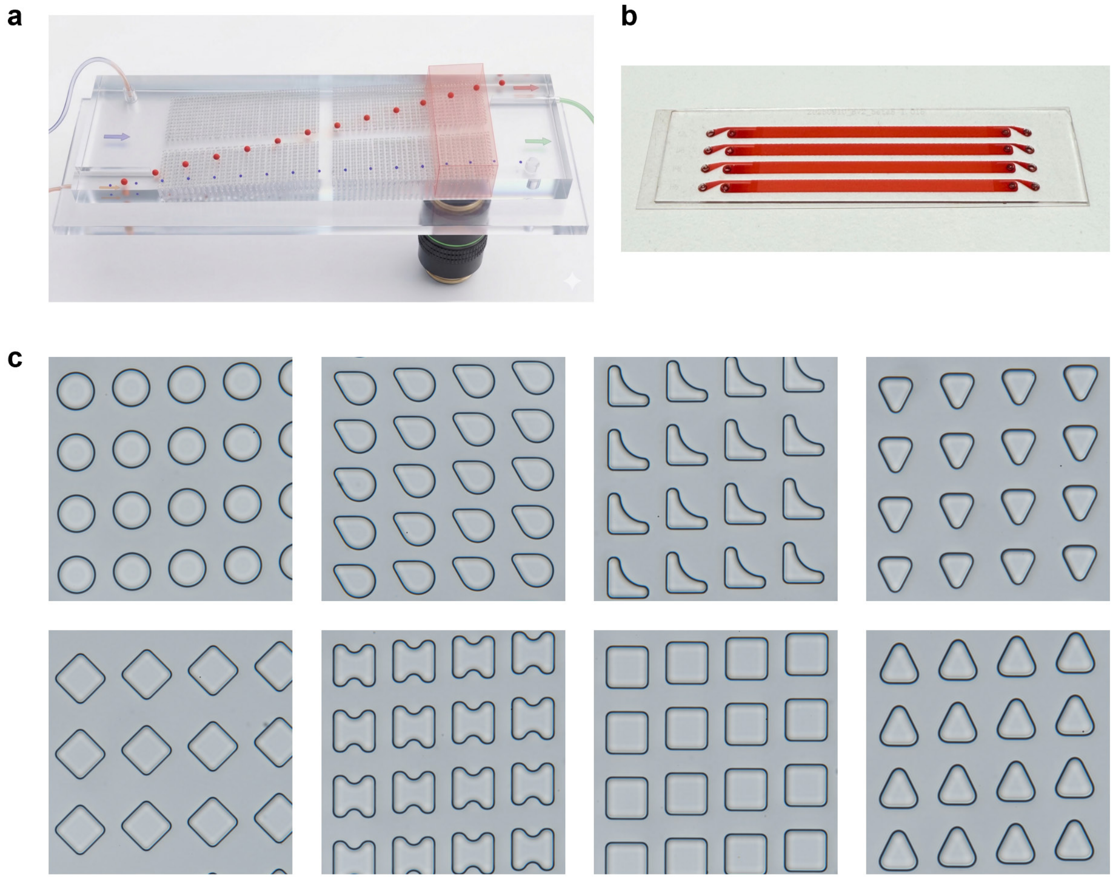

**Figure S1.** Microfluidic deterministic lateral displacement (DLD) devices utilized for the validation of the three-dimensional (3D) fluid-structure interaction (FSI) framework. (**a**) Schematic illustration of the actual experimental trajectories of differently sized particles within a DLD chip, recorded via microscopic imaging. (**b**) Photograph of a physical device with four different geometries. (**c**) Microscopic images showcasing the eight distinct pillar array geometries evaluated throughout this study, including circular, drop-shaped, L-shaped, inverted triangular, diamond, H-shaped, square, and triangular micropillars.

### S2. Statistical determination of critical diameter in DLD devices

To experimentally determine the critical diameters ($D_c$) of DLD arrays, we selected an H-shaped geometry as a representative design. We introduced monodisperse polystyrene microspheres in 0.5 μm size increments into the microfluidic chip and recorded their lateral outlet positions using high-speed microscopic imaging with a particle tracking velocimetry algorithm. By compiling the particle counts at each outlet index into spatial distribution histograms, we identified the effective critical diameter as the specific particle size interval corresponding to the statistical transition from the zigzag to the displacement mode.

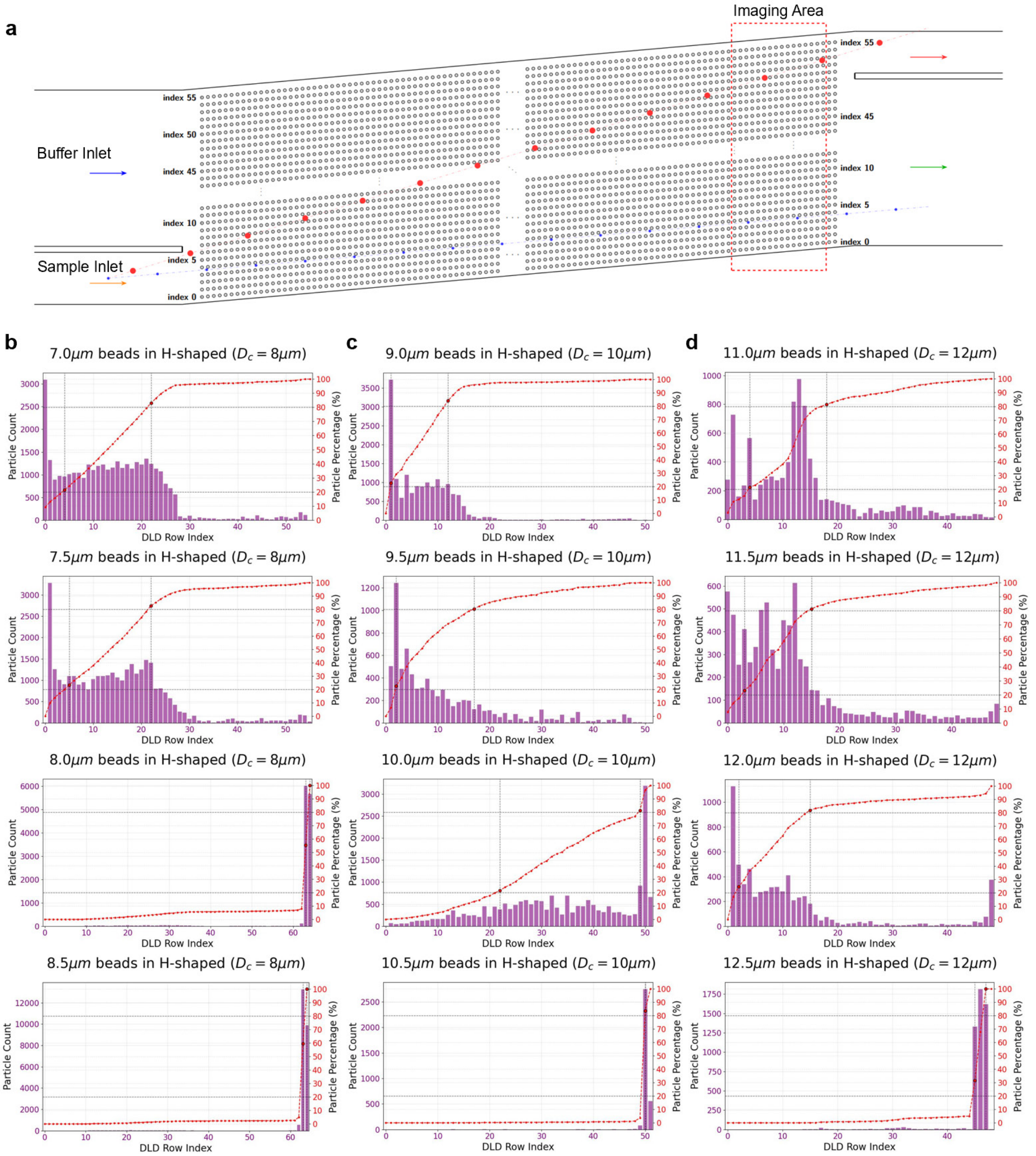

**Figure S2.** Experimental validation of H-shaped pillar DLD arrays across varying critical diameters. (**a**) Schematic illustration of the dual-inlet microfluidic device used for all experimental validations, featuring a sheath flow focusing design. Spatial distribution histograms of particle outlet positions corresponding to three different device designs with theoretical target $D_c$ of (**b**) 8.0 μm, (**c**) 10.0 μm, and (**d**) 12.0 μm, respectively. The bars represent particle counts at each specific outlet index, and the overlaid red dashed curves denote the cumulative particle percentage. By evaluating the transition bandwidths observed in the histograms, the experimentally quantified effective $D_c$ were approximately 8.0 μm, 10.0 μm, and 12.5 μm for the corresponding target structures.

### S3. Enhancement of the computational efficiency

The computational efficiency of the 3D FSI framework is substantially enhanced by implementing a dynamic adaptive mesh. This strategy streamlines spatial discretization by strictly confining high-resolution elements to the fluid-solid interfaces, ensuring that steep hydrodynamic gradients are accurately resolved. This approach enables a reduction in the total number of mesh points by more than 95% compared to a baseline uniform mesh. Consequently, the framework achieves a 91-fold speedup while maintaining accuracy of the high-fidelity simulation.

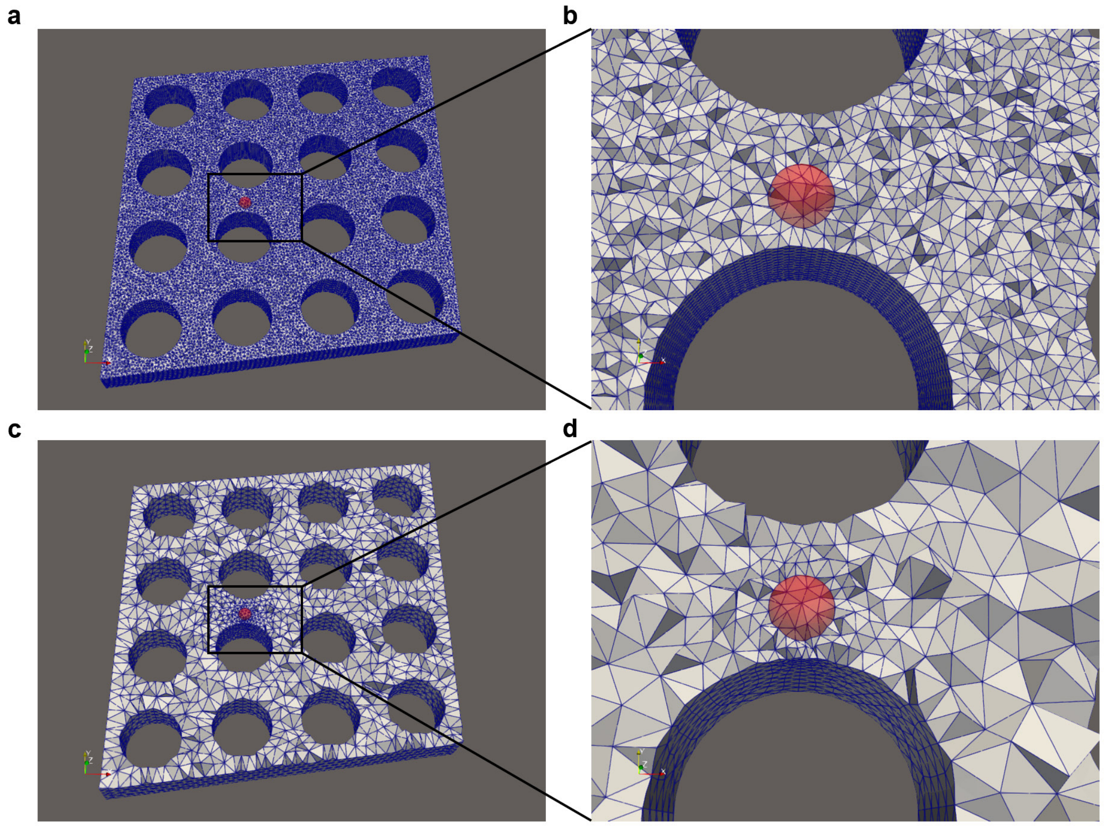


**Figure S3.** Efficiency of dynamic adaptive mesh refinement. (**a**-**b**) Uniform mesh refinement. (**c**-**d**) Dynamic adaptive mesh refinement, localizing high-resolution elements at fluid-solid interfaces. The adaptive strategy preserves computational accuracy while reducing the total mesh points by >95% and achieving a 91-fold speedup.

**Table S1.** Comparison of mesh strategy

| Mesh strategy | Mesh points | Speedup factor |
|---|---|---|
| Uniform mesh refinement | 1,789,565 | 1X |
| Dynamic adaptive mesh refinement | 63,454 | 91X |

Furthermore, the 3D FSI framework accelerates computations by employing high-order discretization schemes in both space and time. For spatial discretization, utilizing second-order isoparametric finite elements provides a 38-fold acceleration compared to a first-order mesh. Similarly, implementing a second-order Runge-Kutta method for temporal integration delivers a 24-fold acceleration over conventional first-order time-stepping techniques. The synergistic integration of these advanced spatial and temporal schemes yields a remarkable 912-fold total speedup relative to a conventional first-order baseline. This combined acceleration approaches three orders of magnitude and serves as the fundamental driver for the transformative computational performance of the framework.

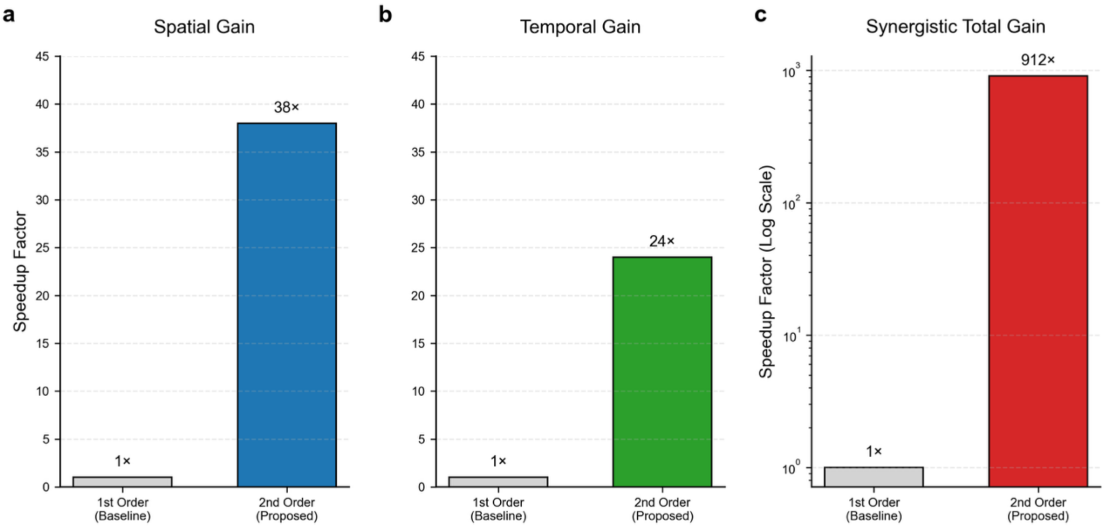


**Figure S4.** High-order discretization efficiency gains. Comparison of speedup factors for (**a**) spatial, (**b**) temporal, and (**c**) synergistic total discretization relative to a first-order baseline. Subfigure (**c**) uses a logarithmic scale to highlight the total acceleration.

### S4. Particle dynamics at the non-periodic device periphery

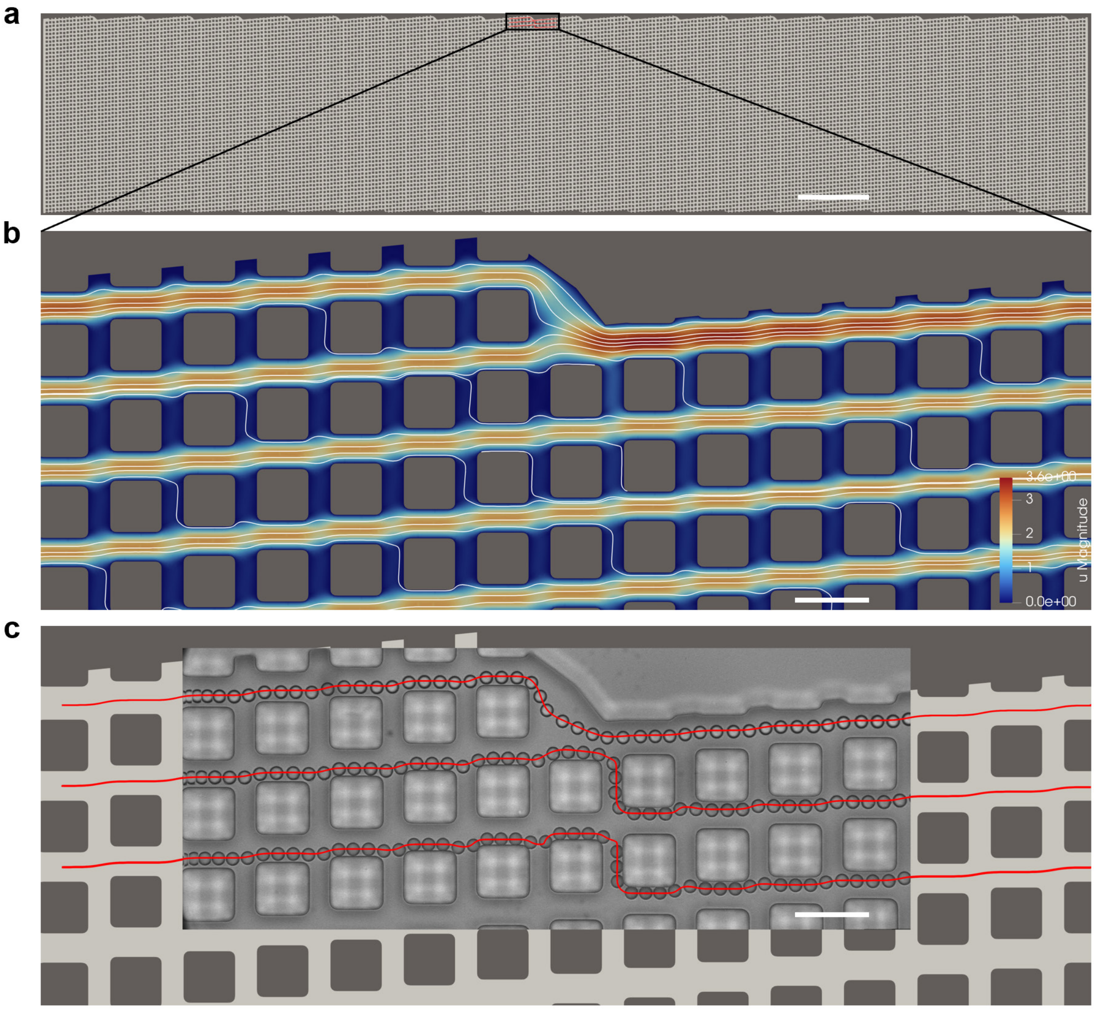


**Figure S5.** Evaluation of experimental and simulated particle dynamics at the non-periodic device periphery. (**a**) Overview of the complete DLD device featuring square micropillars. (**b**) Simulated fluid velocity field near the upper boundary of the device. (**c**) Comparison of experimental and simulated particle trajectories near this upper edge, confirming the framework's capability to accurately resolve boundary layer effects. Scale bars in (**a**), (**b**), (**c**) are 1000 µm, 50 µm and 50 µm, respectively.

**Movie S1. Simulation workflow of the 3D FSI framework**

Movie S1 illustrates the comprehensive simulation workflow of the 3D FSI framework for resolving complex hydrodynamics and particle transport within a full-scale DLD chip. The video illustrates the macroscopic device geometry and introduces the hierarchical and dynamic meshing strategies. To demonstrate the generation of the hierarchical mesh, a background mesh across a vast area is first established to compute the bulk flow field. Subsequently, an intermediate mesh is constructed within a targeted mid-section, followed by the creation of an ultra-fine dynamic mesh in the immediate vicinity of a specific pillar. Crucially, the flow solutions derived from the larger domains serve as accurate boundary conditions for the successively smaller nested regions. The animation then highlights the localized dynamic mesh technique, revealing how the computational grid continuously adapts to track the spatial displacement of the moving particle. Finally, the video presents the results of this fully coupled simulation by displaying the complete 3D trajectory of a single particle navigating the micropillar array from top, side, and front perspectives.

**Movie S2. Comparison of simulated and experimental particle dynamics across diverse array geometries**

Movie S2 sequentially illustrates simulated and experimental particle dynamics across circular, square, drop-shaped, and L-shaped DLD arrays. The video initially visualizes the simulated 3D flow field and streamlines alongside the computed trajectory of a specific particle operating in the zigzag mode. Then the movie presents high-speed videography of physical particles with identical diameters navigating the corresponding fabricated chips. A direct comparison reveals that when launched from identical initial positions, the simulated paths perfectly superimpose onto the actual experimental trajectories.

**Movie S3. Experimental Validation of Simulated Cell Trajectories in a DLD array**

Movie S3 illustrates the simulated and experimental transport dynamics of biological cells within a circular DLD array. The video initially visualizes the simulated 3D flow field and streamlines alongside the computed trajectory of a rigid particle modeled with an equivalent hydraulic diameter. Following this computational demonstration, the movie presents high-speed videography capturing the actual movement of live HT-29 cells navigating the corresponding physical chip. The footage specifically highlights three distinct tracking events originating from varying initial positions. A direct comparison reveals that the physical cellular trajectories broadly superimpose onto the simulated paths. Although minor localized deviations appear near the central pillar regions, the overall transport trend remains highly consistent with the predictive simulations.
